# Metataxonomic analysis of halophilic archaea community in two geothermal springs sources in southern Tunisian Sahara

**DOI:** 10.1101/2023.10.24.563781

**Authors:** Afef Najjari, Khaled Elmnasri, Hanene Cherif, Amel Guesmi, Mouna Mahjoubi, Javier A. Linares-Pastén, Ameur Cherif, Hadda-Imene Ouzari

**Affiliations:** Faculté des Sciences de Tunis, LR03ES03 Laboratoire de Microbiologie et Biomolécules Actives, Université Tunis El Manar, 2092 Tunis, Tunisia; Higher Institute for Biotechnology, University Manouba, BVBGR-LR11ES31, Biotechpole Sidi Thabet, 2020 Ariana, Tunisia; Department of Biotechnology, Faculty of Engineering, Lunds Tekniska Högskola (LTH), Lund University, P. O. Box 124, 22100 Lund, Sweden

**Keywords:** Geothermal springs, Haloarchaea, *Halogranum*, Metataxonomy, Oases

## Abstract

In this study, we assess the phylogenetic diversity of the halophilic archaeal community of biotechnological interest that inhabit in geothermal water in southern Tunisia. These waters are usually used for farming purposes, particularly for irrigation of the oases and greenhouses, due to the limited availability of surface freshwater resources and the arid climate. The samples processed from two separated geothermal sources in southern Tunisia based on the Illumina Miseq sequencing approach. Three samples including water, sediment, and halite-soil crust, were collected from downstream of two geothermal springs of Ksar Ghilane (KGH) and Zaouet Al Aness (ZAN) oases. Results showed that several haloarchaea-related members were identified in geothermal springs. The average taxonomic composition showed that 20 genera out of 33 were shared between the two geothermal sources with uneven distribution, where the *Halogranum* genus is the most represented genus with an abundance of 18.9% and 11.58% for ZAW and KGH, respectively. Several unique site-specific genera were also observed; the case of *Halonotius, Halopelagius, Natronorubrum*, and *Haloarcula* in ZAN, and *Haloprofundus, Halomarina, Halovivax, Haloplanus, Natrinema, Halobium, Natronoarchaeum, Haloterrigena* detected in KGH pool. We noticed that the majority of genus members are usually found in low-salinity ecosystems. The finding of this study suggests that haloarchaea-related members could survive in downstream geothermal sources, which may have resulted from the leaching of salts and minerals through drilling waters creating chemically complex saline systems.

## 1 Introduction

Geothermal springs, also called hot springs, are produced by naturally exothermically heated groundwater. It has been found, over the past years, that geothermal springs harbor a fascinating microbial community capable of withstanding high-temperature conditions. These microbes belonging to Archaea and Bacteria domains are of significant biotechnological importance since they are able to produce valuable biomolecules and thermostable enzymes (Aulitto et al. 2018; Besser et al. 2018; Gallo et al. 2021). For these reasons, a number of studies have been performed that focused on understanding the nature of microbial diversity in such environments based on both culture-independent and culture-dependent approaches in order to obtain a deeper understanding of their taxonomic diversity, adaptive mechanisms, and functional and ecological roles of such microbial communities (Kanokratana et al. 2004; Kato et al. 2009; Sayeh et al. 2010; Schuler et al. 2017; Ghilamicael et al. 2017; Massello et al. 2020).

Assessment of archaeal diversity based on analyses of environmental DNA samples has shown the relative abundance of Crenarchaeota and Thaumarchaeota phyla-related taxa (Burton et al. 2000; Benlloch et al. 2001; Robertson et al. 2005; Huang et al. 2011; Song et al. 2013). Interestingly, some reports have also revealed the presence of halophilic archaea (members of the class Halobacteria) in geothermal springs characterized by their low salinity and high-temperature tolerance (Elshahed et al. 2004a, b; Savage et al. 2007, 2008; Ghilamicael et al. 2017). Some haloarchaea-related members are recovered from hypersaline ecosystems such as salt deposits and solar salterns, and are known to withstand a large range of salinity (0.3%-30% NaCl) (Kim et al. 2018; Najjari et al. 2015, 2021). Moreover, Haloarchaea members were identified in low-salinity environments as brackish waters, sediments and forest soils (Rodriguez-Valera et al. 1979; Munson et al. 1997; Elshahed et al. 2004a, b; Purdy et al. 2004; Savage et al. 2007, 2008; Youssef et al. 2012; Najjari et al. 2015). In fact, Haloarchaea have adapted to survive in their saline environments by employing two main strategies, by uptake and accumulation of high concentrations of inorganic ions such as K^+^ (salt-out strategy) or by synthesis and/or uptake of highly soluble organic solutes that do not interfere with intracellular enzymatic activities and cellular processes (salt-in strategy) (Roberts 2005; Youssef et al. 2014; Becker et al. 2014; Najjari et al. 2015). In addition, Haloarchaea may withstand high temperatures in their natural environment in that some of them have an optimum growth temperature above 50°C, while others retain their enzymatic activity only at high temperatures. Examples include *Natrinema thermophila* (66°C), *Natrinema pellirubrum* (57°C), *Halogeometricum borinquense* (57°C), *Natronolimnobius aegyptiacus* (55°C) and *Natronomonas pharaonis* (56°C) (Robinson et al. 2005; Munawar and Engel 2013; Kim et al. 2018).

Multiple geothermal springs are located in the southern regions of Tunisia. Given the limited availability of surface freshwater resources and the arid climate in southern areas, geothermal waters are typically exploited for oases and greenhouses irrigation (Baccouche 1988; Ben Mohamed 2003). However, the salinity of the geothermal water ranges from 2 to 4.5 g/L and the water is utilized mainly for agriculture purposes can have a negative impact on the irrigated areas due to the accumulation of soluble salts of sodium in soils (Hachicha & Ben Aissa 2014 ; Ben Hassine et al. 1996; Hachicha & Ben Aissa 2014). During an earlier study that examined 16S rRNA gene cloning and sequencing libraries collected from archaeal communities of Tunisian hot springs, a number of haloarchaeal clones were revealed (Sayeh et al. 2010). To our understanding of haloarchaeal communities in such areas, we used a specific 16S rRNA gene-based amplicon sequencing approach to investigate haloarchaeal communities in sediments, halite soil crusts, and waters collected from two geothermal springs located in the southern part of Tunisia. As such, this study has deepened our understanding of the haloarchaeal diversity of southern Tunisian hot springs.

## 2 Material and methods

### 2.1 Sampling and sites description

Samples were collected in December 2019 from two geothermal hot springs sources in the Kebili governorate in southwestern Tunisia (see (i) and (ii) below for details of locations). The area is characterized by an arid desert climate. Salinities were measured using a hand-held SW series VistaVision refractometer (VWR, Radnor, PA). Temperature and pH were recorded on site, and*≈*100g/1000mL was sampled into sterile containers, placed on ice, and transported to the laboratory, where the samples were kept frozen (-20°C) until DNA extraction.

1. The first source is the deep geothermal drill of Zaouiet El Anes (ZAN) (33047’30.69”N; 8047’22.03”E; 2358m deep) in Souk Lahad delegation (Fig. 1a,b). The pH of drilling waters is neutral (7), the salinity content of 4.5 g/L, and the temperature ranges from 45°C downstream of the source, to 71°C at the top of the geothermal source. Water sources are used for irrigation of oasis and bathing purposes. Samples identified as S35 (sediment), S33 (water), and S36 (halite-crust soil) were taken from downstream of the geothermal source where the temperature of water is about 45°C.
2. The second source is the geothermal drill site of Ksar Ghilane oasis (KGH) (32059’18.04”N; 9038’22.25”E; 580m deep) located in the middle of sand dunes of Douz delegation (Fig. 1b). As with the first site, water sources of KGH are used for irrigation of oasis and bathing purposes. The water of the source is characterized by a temperature of 42°C, pH 7.5, and salt content of 4,5 g/L. From this source, three samples were taken from from the edge of the KGH water pool (Fig. 1a,c) and were identified as S20 (sediment), S23 (water), and S24 (halite crust soil).

**Figure 1:**
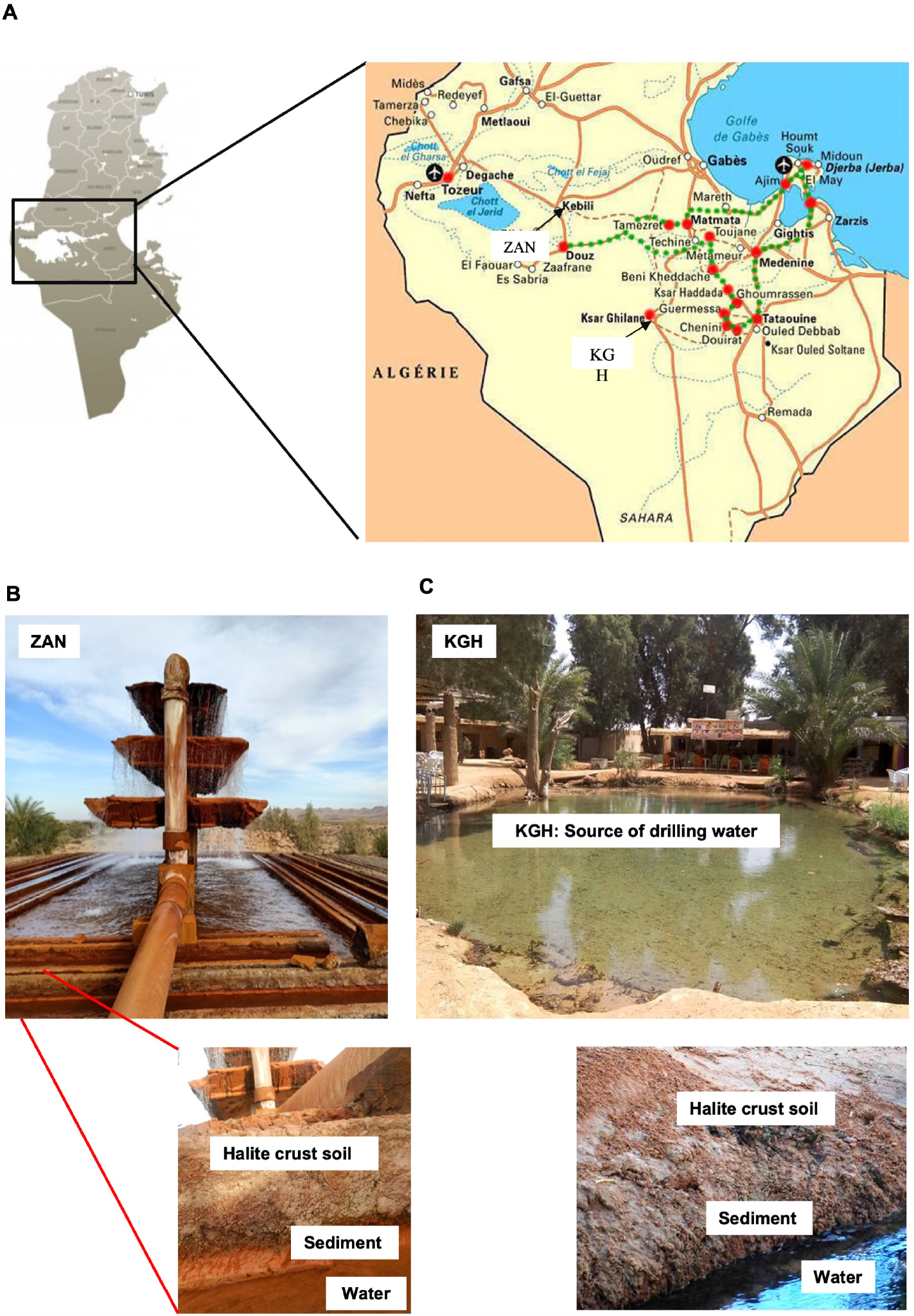
(a): Geothermal hot springs sources localization in the Tunisian map; (b): geothermal source in ZawietAl Aness (ZAN), (c); Pool of geothermal water in KsarGhilane..

### 2.2 Culture-independent assessment

#### 2.2.1 DNA extraction, PCR amplification, sequencing, and analysis

DNA was extracted using a PowerSoil DNA extraction kit (MoBio, Carlsbad, CA) following the manufacturer’s instructions and quantified using a Qubit fluorometer (Life Technologies, Grand Island, NY). Halobacteria community structure was targeted using Halobacteria-specific primers (287F and 589R) (Najjari et al. 2015). PCR analysis was performed in 50 *μ*L reaction mixtures that contained 2-5 *μ*L of the extracted DNA, 1*×*PCR buffer (Fermentas), 2.5 mM MgSO_4_, a 0.2 mM deoxyribonucleoside triphosphate (dNTP) mixture, 0.5 U of Taq DNA polymerse (Fermentas), and a 10 *μ*M concentration of each of the forward and reverse primers. PCR was carried out according to the following protocol: initial denaturation at 95°C for 5 min, followed by 35 cycles of denaturation at 95°C for 45 s, annealing at 55°C for 1 min, and elongation at 72°C for 1 min. A final elongation step at 72°C for 10 min was included. Sequencing libraries were prepared using Nextera XT kit according to Illumina Supporting Guide (Klindworth et al. 2012). Libraries were then multiplexed and sequenced on the MiSeq paired-end Illumina platform (Illumina, San Diego, CA, USA).

#### 2.2.2 Bioinformatics analysis for abundance and taxonomic assessment

The DADA2 (Divisive Amplicon Denoising Algorithm) microbiome pipeline v1.22 (Callahan et al. 2016) embedded in R software v4.2.0 (R Core Team, 2021) was used for the paired-end reads the analysis. Sequences were classified into operational taxonomic units (OTUs at 0.03 cutoff) using SILVA non-redundant v138 training set (Quast et al. 2012). Rarefaction curves were computed. The number of OTUs at the genus level was assessed to describe alpha diversity between the different groups. Alpha diversity indices corresponding to variables Good’s coverage, Shannon’s diversity index (H), Chao 1, and Simpson were conducted using Mothur software (Schloss et al. 2009). The Shannon and Simpson *α*-diversity indices were applied to estimate the diversity for each group using the Wilcoxon rank-sum test (Yoon et al., 2017). Beta diversity was calculated with Bray-Curtis distances based on the taxonomic abundance profiles. Permutational multivariate analysis of variance (PERMANOVA) was applied to measure the statistical significance of *β*-diversity (Yoon et al., 2017). The significance level applied for all statistical tests was 5% (*p <* 0.05).

Data availability. The obtained raw sequences were submitted to the National Center for Biotechnology Information (NCBI) as Sequence Read Archive (SRA) under accession numbers as follows S33 (SRR23374283), S36(SRR23423409), S35 (SRR23423408), S23 (SRR23423493), S20 (SRR23423410) and S24 (SRR23423411).

## 3 Results and Discussion

### 3.1 Physicochemical parameters of water and sediment samples

Salinity, temperature, pH, conductivity, dry residue, and the concentrations of ions such as Ca^2+^, Mg^2+^, Na^+^, K^+^, SO_4_^2−^, Cl^−^, NO_3_^−^, and HCO_3_^−^ were determined from the oases samples, KGH and ZAN (Table 1). The salinity values of both water and sediment samples are within the range expected for oases, with slightly higher values for KGH compared to ZAN. The temperature values are also within the range expected for arid environments, with KGH having a somewhat higher temperature than ZAN. The pH values are slightly alkaline for both oases, with ZAN having a slightly higher pH than KGH. The conductivity values are higher for ZAN compared to KGH, although KGH has higher concentrations of the ions analyzed. Indeed, the concentrations of Ca^2+^, Mg^2+^, Na^+^, Cl^−^, and SO_4_^2−^ are higher in KGH compared to ZAN, while the concentration of K^+^ is higher in ZAN compared to KGH. Both oases have low concentrations of NO_3_^−^. The dry residue values for both water and sediment samples are comparable for both oases, indicating a similar level of dissolved solids. Overall, the physicochemical characteristics of the two oases show some differences in the ion composition of their water and sediment samples. These differences may be due to variations in geological, hydrological, and biological factors between the two oases.

**Table 1:** Chemical composition of samples collected from KGH and ZAN sites.

| Parameter | KGH |  | ZAN |  |
| --- | --- | --- | --- | --- |
|  | water | sediment | water | sediment |
| Salinity (g/L) | 5.7 | 4.7 | 4 | 4.2 |
| Temperature (°C) | 43.2 |  | 38 |  |
| pH | 7.6 | 7.6 | 7.8 |  |
| Conductivity (mS/cm) | 5.7 | 6 | 6.74 | 6.13 |
| Dry residue (mg/L) | 3853 | 3770 | 3853 | 3750 |
| Ca<sup>2+</sup> (mg/L) | 483 | 320 | 337.6 | 320 |
| Mg<sup>2+</sup> (mg/L) | 237.6 | 226.8 | 171.9 | 222 |
| Na<sup>+</sup> (mg/L) | 849.3 | 850.03 | 386.4 | 333.5 |
| K<sup>+</sup> (mg/L) | 19 | 20.1 | 51.48 | 46.02 |
| SO<sub>4</sub><sup>2-</sup> (mg/L) | 1561 | 1102.02 | 996 | 859.2 |
| Cl<sup>-</sup> (mg/L) | 1517.7 | 1429.5 | 709 | 1057.9 |
| NO<sub>3</sub><sup>-</sup> (mg/L) | 0.1 | No detected | 0.4 | 6.1 |
| HCO<sub>3</sub><sup>-</sup> (mg/L) | 384 | 393.7 | 115 | 384.4 |

### 3.2 Diversity and richness estimates

Diversity indices of the archaeal community are illustrated in Table 2. The obtained sequence data represented a diverse haloarchaeal community of 622 and 371 OTUs for KGH and ZAN samples’ respectively (3% divergence). Rarefaction curves and good coverage with values *>*98% for all samples indicate an overall excellent OTU coverage afforded by the deep sequencing (Fig. S1). Species richness and diversity index values among KGH and ZAN sets are compared and statistically tested (Fig. 2). Results showed that the species richness of KGH samples was higher than ZAN as measured by Chao 1 (*p* = 0.04). Indeed, the diversity index, as calculated using the Shannon, Simpson, and Bray-Curtis indices, showed also a significantly difference between the two sets at the genus level (Fig. 2). These differences may be due to (i) the chemical composition of waters, where water from KGH site is highly mineralized compared to ZAN (Table 1).

**Table 2:** Alpha-diversity of the haloarchaeal communities associated with KGH and ZAN samples.

| Sample name | Target reads | OTUs | CHAO | Shannon | Simpson | Good's coverage of library (%) |
| --- | --- | --- | --- | --- | --- | --- |
| S23_ground water | 15325 | 953 | 954.17 | 5.12 | 0.02 | 99.82 |
| S20_sediment | 6518 | 296 | 303.56 | 4.57 | 0.02 | 99.57 |
| S24_halite crust soil | 2909 | 233 | 252.02 | 4.45 | 0.02 | 98.62 |
| S33_ground water | 1694 | 127 | 135.05 | 3.98 | 0.04 | 98.94 |
| S35_sediment | 1242 | 73 | 77.59 | 2.88 | 0.12 | 98.95 |
| S36_halite crust soil | 1379 | 171 | 177.66 | 4.26 | 0.04 | 98.33 |

**Figure 2:**
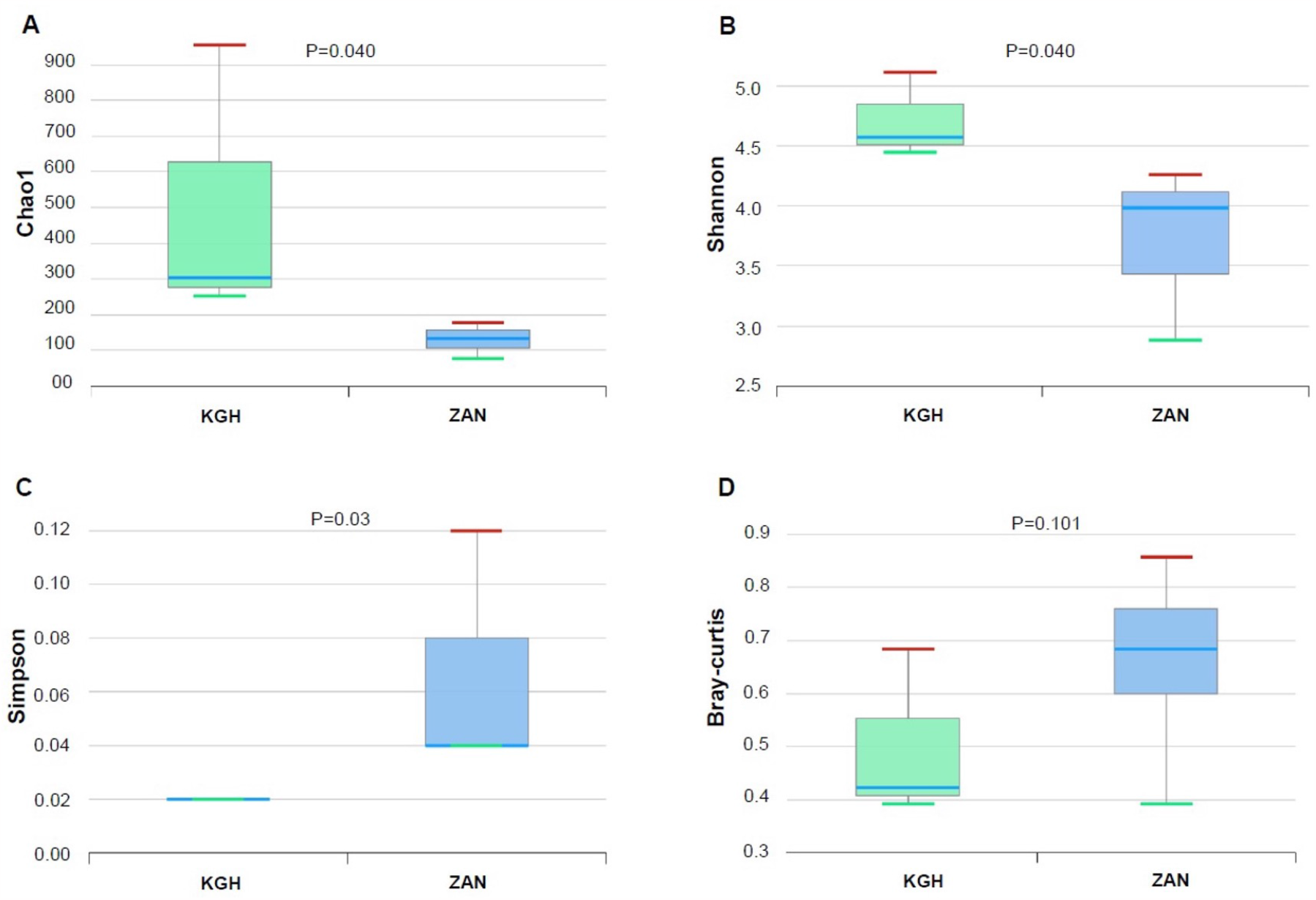
Comparison of species richness and *α*-, *β*-diversities in haloarchaeal taxonomic profile between ZAN and KGH samples sets by (A). Chao1, (B). Shannon, (C). Simpson, and (D). BrayCurtis indices. The Shannon and Simpson *α*-diversity indices were applied to estimate the diversity for each group using the Wilcoxon rank-sum test. Beta diversity was calculated with Bray-Curtis distances based on the taxonomic abundance profiles. Permutational multivariate analysis of variance (PERMANOVA) was applied to measure the statistical significances of *β*-diversity.

### 3.3 Taxonomic distribution and phylogenetic assignment of Haloarchaeal members

The genera distribution found in samples from geothermal spring sources (ZAN and KGH), assessed based on Halobacteria-specific gene amplifications, are shown in Table 3.

**Table 3:**
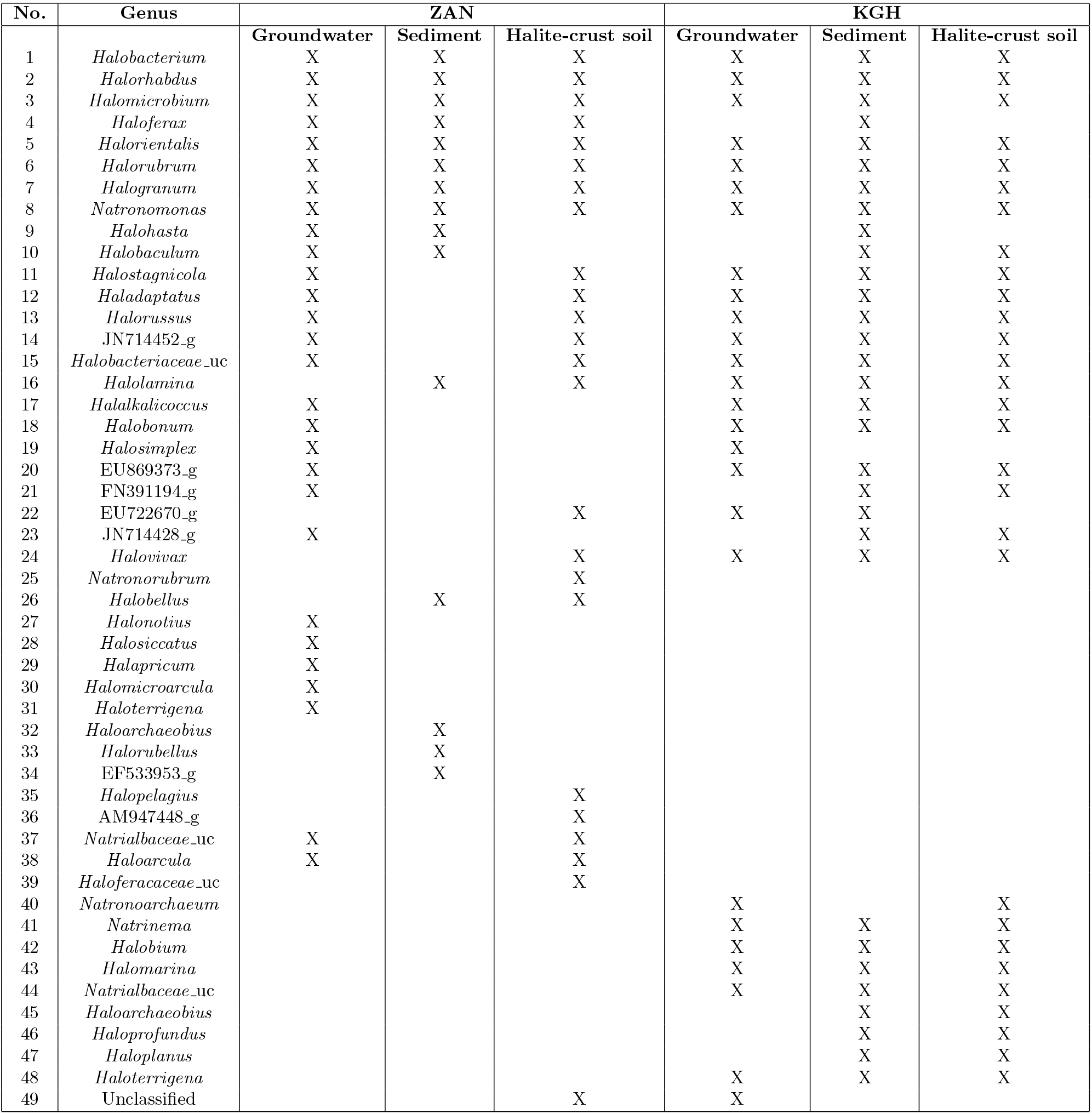
Genera distribution across the samples.

#### 3.3.1 ZAN samples

A total number of 4315 valid reads amplicons was produced with an average length of 280 bp across 3 samples ranging from (150 bp to 300 bp) (Table 2). Taxonomic level distribution showed that OTUs of all samples (*n* = 3) were affiliated to the Euryarchaeota phylum and Halobacteria class. Three dominant orders were identified, with Haloferacales being the most abundant members in the halite-crust soil (84.62%), followed by Halobacteriales, which was similarly represented in sediment and groundwater samples (59%), then Natrialbales-related members were identified in ground water and sediment samples only (10.29 and 5.84% respectively). At the genus level, a total of 27 genera were identified out of 51 halophilic archaea genera presently described (Parte et al. 2020) and distributed as follows:

- In the groundwater sample (S33), a total of 26 genera have been identified with a net predominance of Halohasta (16.64%) (Fig. 3). The remaining genera members were less than 9% in relative abundance. The genus *Halohasta*, an haloalkaliphilic archaeon, was first identified in 2012 and includes two species, *Halohasta litorea*, and *Halohasta litchfieldiae*, isolated from a brine sample from an aquaculture farm in Daliang, China, and a surface water sample from Deep Lake, Antarctica, respectively (Mou et al. 2012).
- Within the halite soil crust sample (S36), 20 genera were identified with the occurrence of *Halorientalis* genus members (22.19%) (Fig. 3). The remaining genera comprise less than 6% of the relative abundances. The genus *Halorientalis* was proposed by (Cui et al. 2011) and actually includes five species, *Hos. brevis, Hos. hydrocarbonoclasticus, Hos. pallida, Hos. persicus*, and *Hos. regularis* (Parte et al. 2020). These species have been isolated from saline environments (Durán-Viseras et al. 2019), and recently *Halorientalis* was identified in freshwater from the Liaohe estuary (China) (Shi et al. 2021).
- In the sediment sample (S35), 13 genera were identified with the occurrence of sequences affiliated with *Halogranum* genus members being the most abundant (48.12%) (Table 3, Fig. 3). The relative abundance of remaining genera does not reach 5% of total classified reads. In fact, the occurrence of the Halogranum genus in low-salinity habitats was previously reported (Youssef et al. 2012; Najjari et al. 2015). Currently, four *Halogranum* species are described including *H. amylolyticum, H. gelatinilyticum, H. rubrum*, and *H. solarium* (Cui et al. 2010; Cui et al. 2011). *Hgn. rubrum* and *Hgn. gelatinilyticum* species are the most abundant in the sediment of ZAN (25% and 20% respectively). Unaffiliated species sequences with low abundance (1%) were also detected. *Hgn. rubrum* and *Hgn. gelatinilyticum* species have been previously isolated from marine solar salterns and evaporitic salt crystals collected along the seashore of Namhae, South Korea (Cui et al. 2010, 2011; Kim et al. 2011).

**Figure 3:**
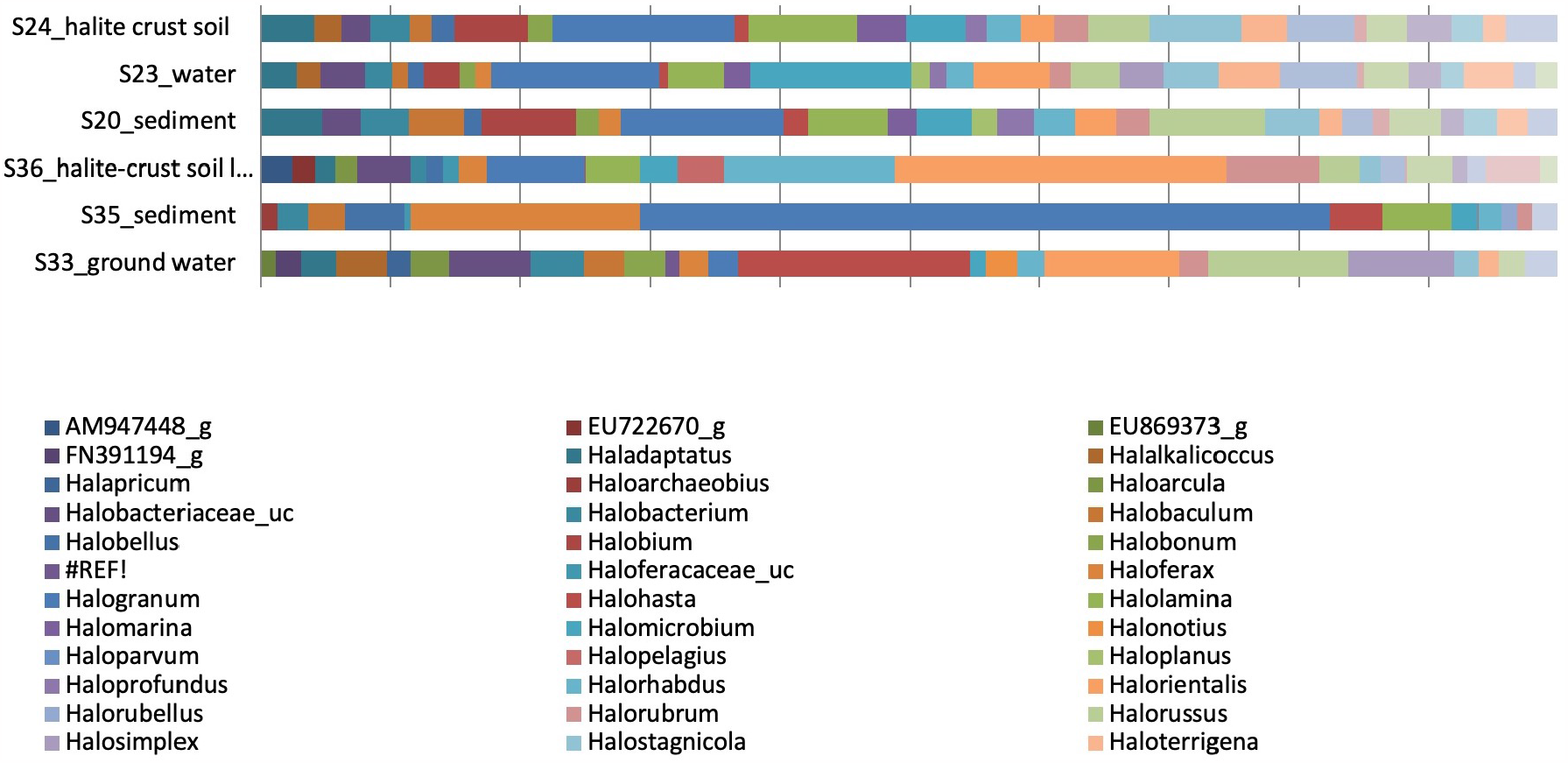
Bar plots showing the genus taxonomic abundance distributed across the samples. Taxa with low abundance (*<*1%) was eliminated

The nature and prevalent salinities in these samples suggest the capability of the genus to survive in environments of varying salinities (Youssef et al. 2014; Najjari et al. 2015; Shi et al.2021). It is noteworthy that eight main genera were common to all samples, comprising *Halobacterium, Halorhabdus, Halomicrobium, Haloferax, Halorientalis, Halorubrum, Halogranum*, and *Natronomonas*, the majority of which are widespread in nature, ranging from salt-saturated habitats to low-saline environments, as well as from hot spring sources (Purdy et al. 2004; Elshahed et al. 2004a, b; Sayeh et al. 2010; Youssef et al. 2014; Najjari et al. 2015, 2021).

A further comparison of the haloarchaea genera members indicated that three samples (water, halite crust soil, and sediment) shared 8 common genera including *Halobacterium, Halorhabus, Halomicrobium, Haloferax, Halorientalis, Halorubrum Halogranum* and *Natronomonas* (Fig. S2, Table 3). Genera unique to individual samples were also observed including *Halalkalicoccus, Halobonum, Halonotius, Halosiccatus, Halapricum, Halomicroarcula, Haloterrigena, Halosimplex* and uncultured *Haloarchaeaon* (FN391194, EU869373, JN714428) which were only detected in the water sample, while the genera *Halopelagius, Natronorubrum*, and *Halovivax* (plus some unclassified and uncultured genera) were detected only in the halite soil crust (Fig. S2, Table 3). Finally, *Haloarchaeobius, Halorubellus*, and an uncultured *Haloarachaeaon* were specific to the sediment samples (Fig. S2, Table 3).

#### 3.3.2 KGH source

A total number of 24,752 valid reads amplicons was produced with an average length of 282 bp across 3 samples ranging from 100 bp to 300 bp (Table 1). Similar to the ZAN pool, the OTUs of all datasets were affiliated with the Euryarchaeota phylum and Halobacteria class. Three Orders were identified within all samples, represented by Halobacteriale which were the most abundant with relative abundances of 44.93, 48.52, and 39.70% for S20, S23, and S24 respectively. The Haloferacales represent 41% of the Halobacteria class in the sediment, 37.74% in the halite soil crust, and 28.84% in water. In the position of least abundance, we find Natrialbales members that represent 13.11% of sediment, 21.12% of groundwater, and 22% of halite soil crust Halobacteria.

At the genus level, a total of 27 genera were identified in all samples with unequal distribution (Fig. 4). Datasets of three samples contain one main genus-related member, the Halogranum with a relative abundance of*≈*12%. Indeed, a total of 22 genera (*Halomarina, Halostagnicola, Halolamina, Halalkalicoccus, Halobacterium, Halobonum, Halorhabdus, Halomicrobium, Halorientalis, Haladaptatus, Halorussus, Halovivax, Halorubrum, Natrinema, Halogranum, Halobium, Natronomonas*, unclassified-Halobacteriaceae related members, *Haloterrigena*, and unclassified *Natrialbaceae* related members) are common in all KGH samples (Fig. S2, Table 3). Other specific genera were detected within the halite-crust soil sample (Fig. S3, Table 3).

**Figure 4:**
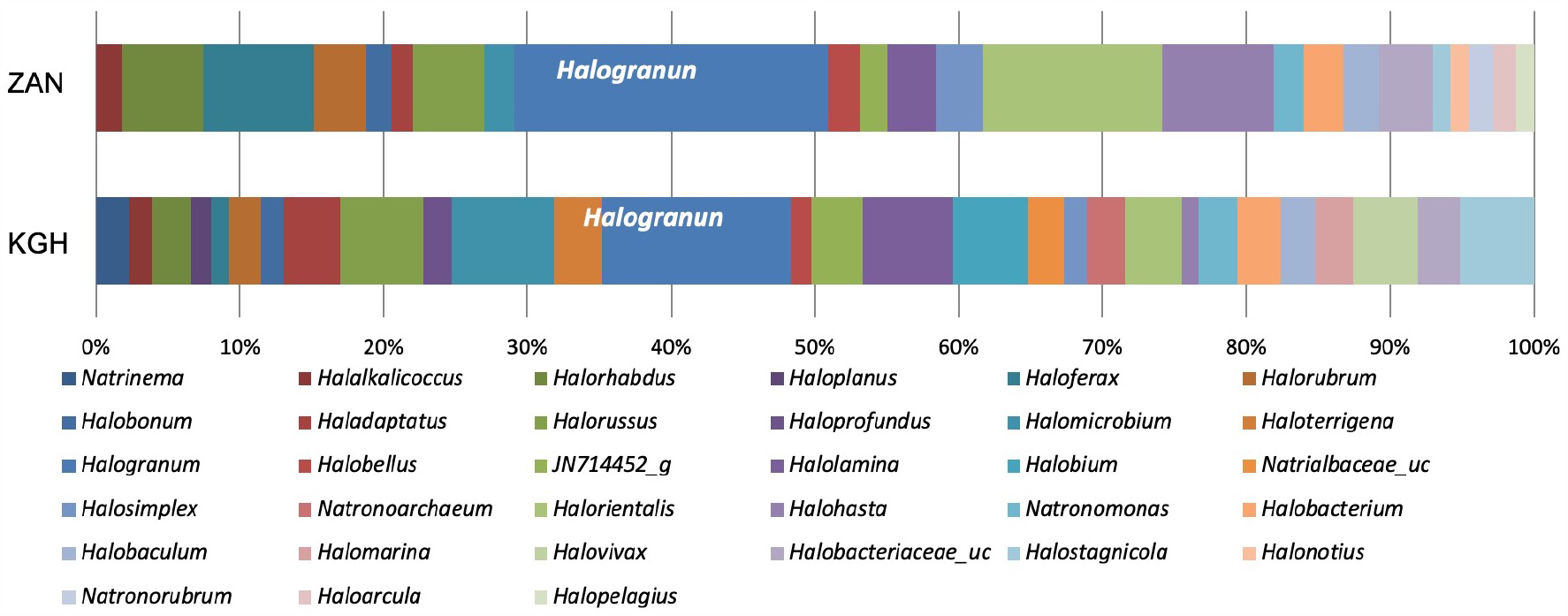
Bar plots showing the average taxonomic composition of genus members across the ZAN and KGH sites. Taxa with low abundance (*<*1¡1%) was eliminated.

### 3.4 Average taxonomic composition of two geothermal sources

The average taxonomic compositions of shared and unique archaeal taxa between the two datasets were assessed (Fig. 4). Results showed noticeable differences in haloarchaeal communities’ distribution between the two geothermal sites.

Among the 33 detected genera, 20 were encountered in the two geothermal sources investigated, with uneven distribution (Fig. S4, Table 3). It’s worth noting that *Halogarnum*-related sequences are the most represented across the two sites with an abundance of 18.9% and 11.58% for ZAN and KGH sites respectively (Fig. 5). The remaining genera including *Halostagnicola, Halolamina, Halalkalicoccus, Halobacterium, Halobonum, Halorubrum, Halohasta, Halobaculum, Halorhabdus, Halomicrobium*, JN714452 g, *Haloferax, Halorientalis, Natronomonas, Haladaptatus, Halobacteriaceae* uc, *Halorussus, Halosimplex*, and *Halobellus* represent less than 10% of the haloarchaeal community. The genera *Halonotius, Halopelagius, Natronorubrum*, and *Haloarcula* are detected only in ZAN samples, while eight genera were detected only in KGH samples, comprising *Haloprofundus, Halomarina, Halovivax, Haloplanus, Natrinema, Halobium, Natronoarchaeum, Haloterrigena*, as well as several unclassified *Natrialbaceae* members. Almost all of the above genera were previously identified in low-salinity environments. In a survey of a low-salt, sulfide, and sulfur-rich spring in southwestern Oklahoma, Elshahed, and collaborators isolated halophilic archaeal clones assigned to *Halogeometricum, Natronomonas, Halococcus*, and *Haloferax* genus members (Elshahed et al. 2004a). The salinity of brine and sediment samples from which these archaea are isolated ranges between 0.7 to 1%. Similarly, *Haloferax* and *Halogeometricum* members have been isolated from low-salinity estuary sediments (Purdy et al. 2004). Haloarcula and *Halobacterium* members have been isolated from low-salinity coastal sediments and brine of Goa (Braganca and Furtado 2009).

**Figure 5:**
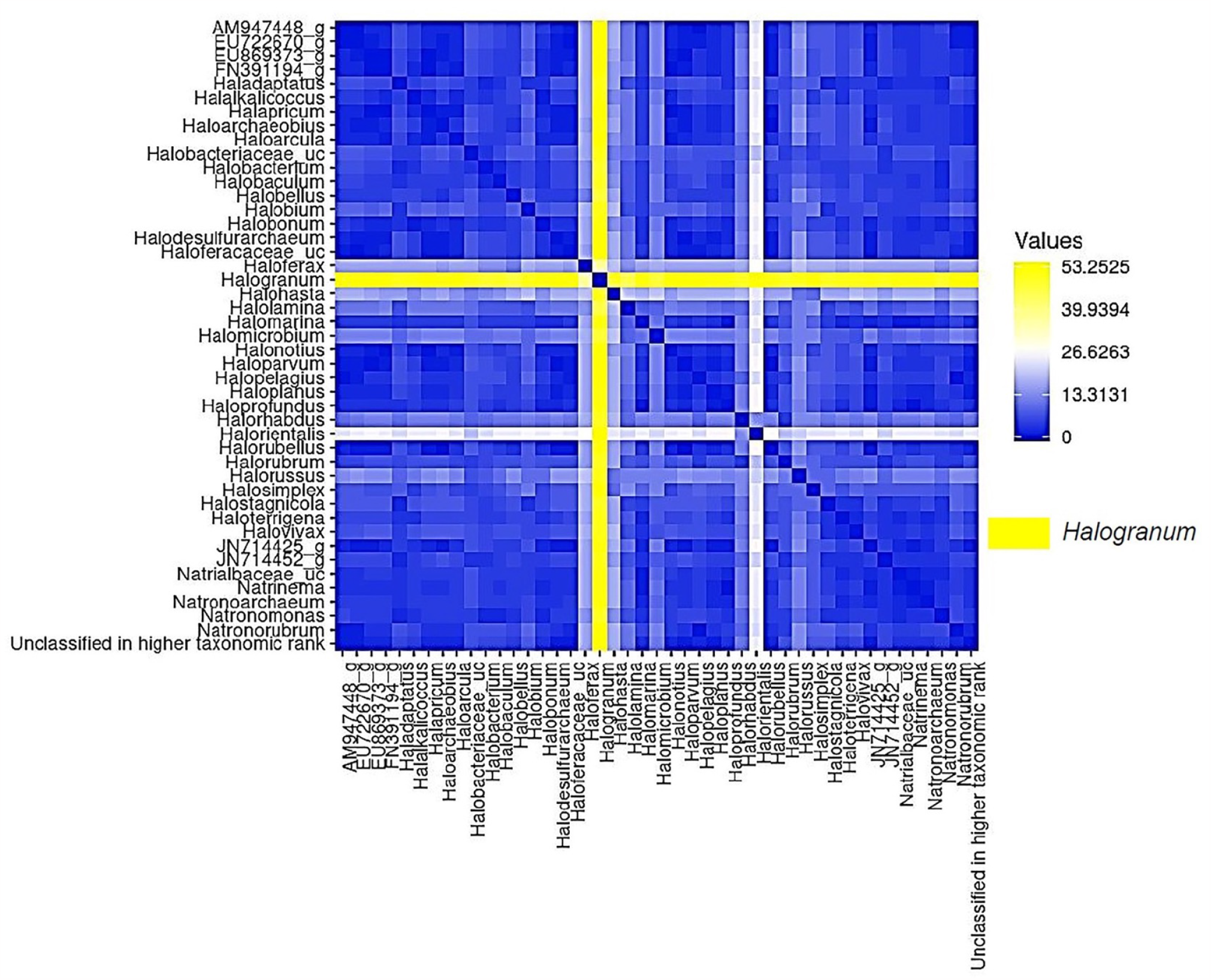
Hierarchical clustering dendrogram based on a distance matrix of average taxonomic compostion at genus level in KGH and ZAN sites showing the abundance of *Halogranum* genus members comparing to other genera.

Recently, Kimbrel et al (2018) have illustrated the identification of halophilic archaea in brine and sediment (7.5% NaCl) collected from a solar salt pond. Some Haloarchaea clones have also been detected in another Tunisian study, which used 16S rRNA gene libraries to evaluate a Mahassen geothermal source in southern Tunisia (Sayeh et al. 2010). It is noteworthy that Mahassen, ZAN, and KGH springs form part of the Continental Intercalary (CI) aquifer of southern Algeria and Tunisia (Abid et al. 2009).

Collectively, the results suggest that Halogranum–related sequences are the most abundant genus in our samples. To our knowledge, no studies describe the survival mechanisms responsible for enabling such microorganisms to thrive under conditions of low salinity and high temperature simultaneously. Possibly, the existence of haloarchaea-related members may be the result of water diffusing to the borders of the hot spring water channel, where its evaporation may induce a reduction in humidity and an increase in salinity in the top layers of the soil, thus providing a suitable environment for halophilic archaea. Further work is recommended to test such a hypothesis. To adapt to the low salt environment, halophilic archaea use a variety of organic solutes to counter external osmotic pressure including amino acids and derivatives, polyols, and derivatives, sugars (trehalose), betaines and ectoine (Youssef et al. 2014; Najjari et al. 2015; Najjari et al. 2023). Indeed, a correlation between intracellular accumulation of compatible solutes and growth at supraoptimal temperatures has been observed in halotolerant or slightly halophilic (hyper) thermophiles, indicating that these solutes could act in a thermostabilization function (Hensel and König 1988; Martins and Santos 1995; Ramos et al. 1997; Da Costa et al. 1998; Pais et al. 2005).

## 4 Conclusion

Summarize the key points of youHere we investigate the haloarchaeal diversity in samples (halite-crust soil, ground waters, and sediments) collected from two geothermal springs sources in southern Tunisian Sahara (KGH and ZNA), based on Illumina Miseq sequencing targeting V3-V4 regions. The results showed a distinction in taxonomic distribution between samples of the sample site and also between two sites (ZNA and KGH). The amalgamation of all results revealed the dominance of *Halogranum*-related sequences in two sites. Indeed, some of OTUs were only detected in KGH, like *Haloprofundus, Halomarina, Halovivax, Haloplanus, Natrinema, Halobium, Natronoarchaeum, Haloterrigena*, while others were related to ZAN only citing *Halonotus, Halopelagius, Natronorubrum*, and *Haloarcula*. These differences may be due to the difference in the chemical composition of samples, their salinities, and the flow velocities of running waters. Further studies based on the whole genome method (WGS) could enhance our understanding of the mechanisms adopted by haloarchaea members to colonize geothermal springs sources.r research.

## Acknowledgments

This work was supported by the National Science Foundation Microbial Observatories Program (grant EF0801858), and by the BIODESERT Project (European Community,,s Seventh Framework Program CSA, EU FP7-CSA-SA REGPOT-2008-2; grant 245746).

## Author contributions

AN, AC, JAL-P and HO designed, and wrote the manuscript. AN and KE conducted experiment. AN, KE, HC, AG and MM contribute to write methodology and AN did bioinformatics and statistical analyses. All authors read and approved the manuscript.

## Notes

### Competing Interest Statement

The authors have declared no competing interest.

