## Supplementary Material for "Metataxonomic analysis of halophilic archaea community in two geothermal springs sources in southern Tunisian Sahara"

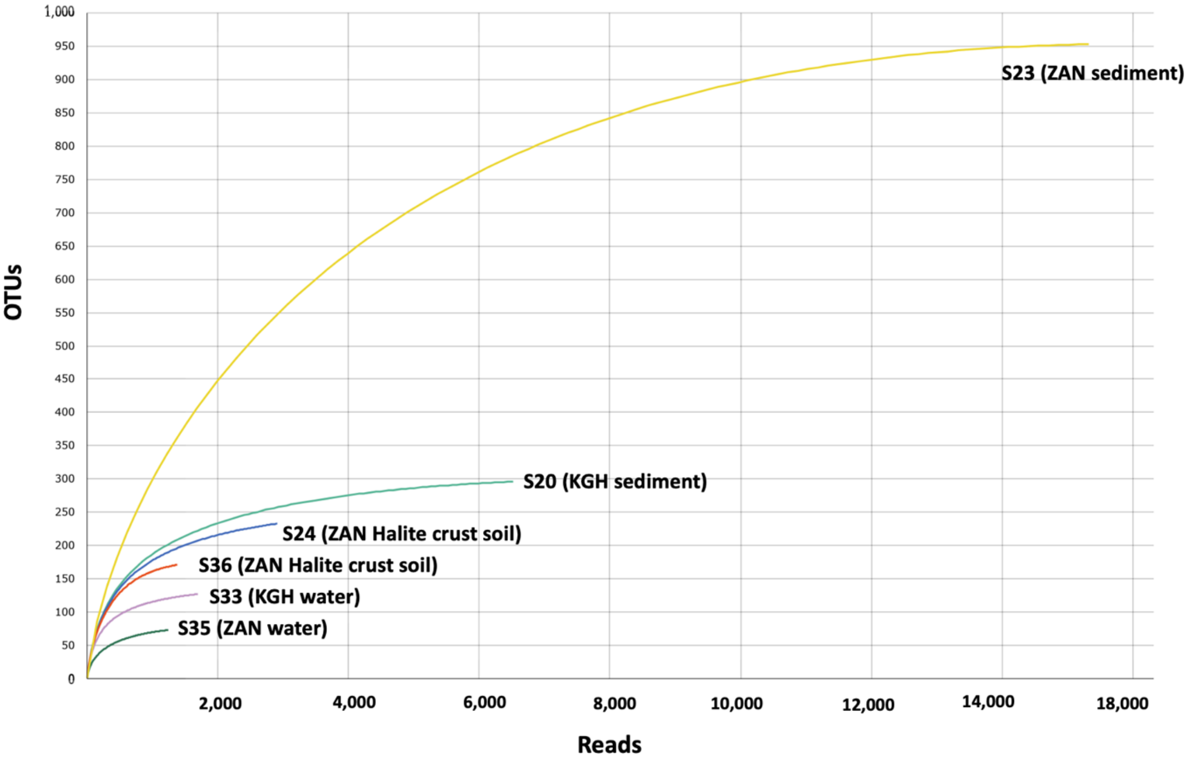


**Fig. S1** Genus-level rarefaction curves of samples based on the 16S rDNA sequences. OTUs are operation taxonomic units.

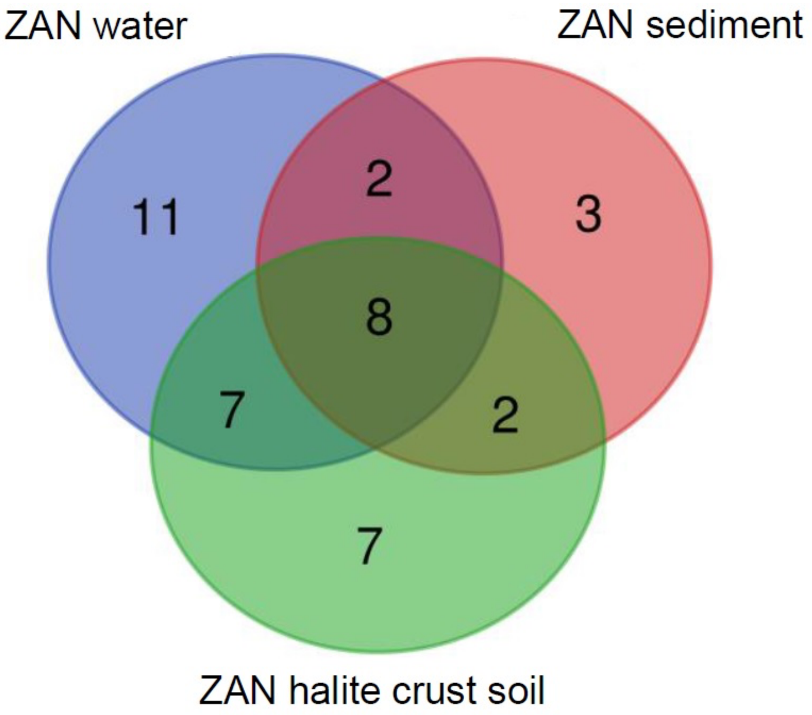


**Fig. S2:** Diagram representing the distribution of common and unique OTUs (3% distance level) across samples of ZAN source.

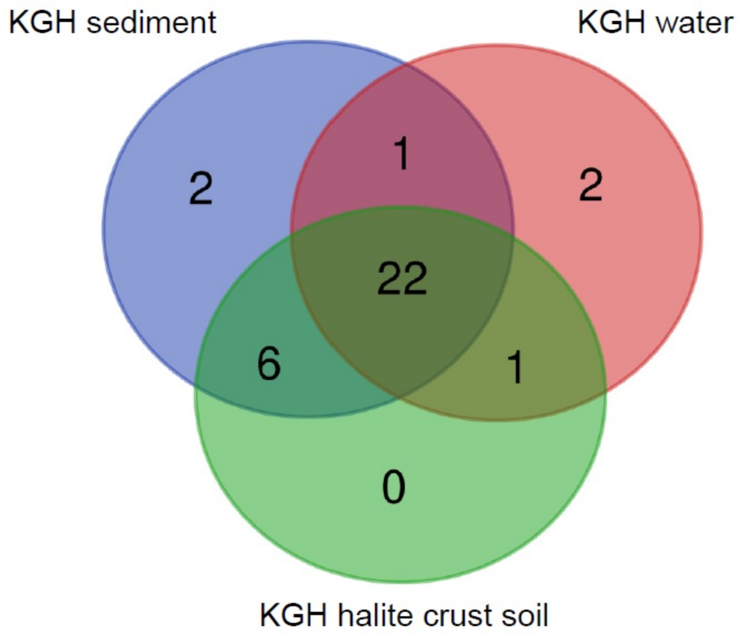


**Figure S3:** Diagram representing the distribution of common and unique OTUs (3% distance level) across samples of KGH source.

**Supplementary Material**

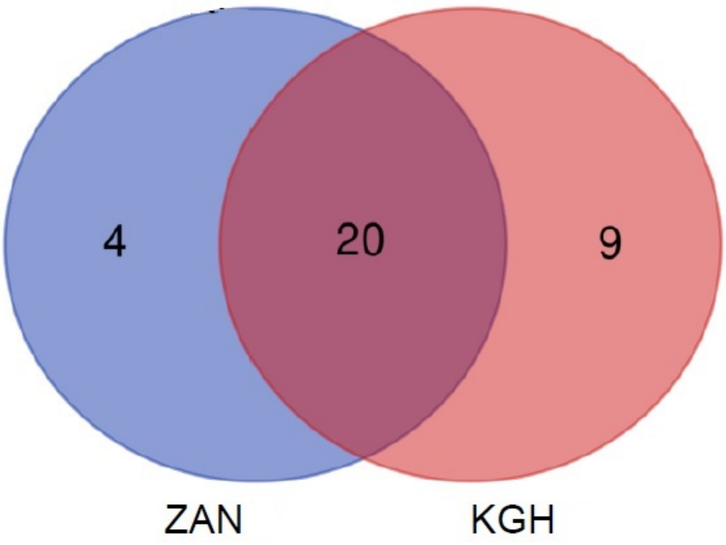


**Figure S4:** Diagram representing the distribution of common and unique OTUs (3% distance level) across KGH and ZAN geothermal spring sources,
